# Computational studies reveal structural characterization and novel families of *Puccinia striiformis* f. sp. *tritici* effectors

**DOI:** 10.1101/2024.09.12.612600

**Authors:** Raheel Asghar, Nan Wu, Noman Ali, Yulei Wang, Mahinur Akkaya

## Abstract

Understanding the biological functions of *Puccinia striiformis* f. sp. *tritici* (*Pst*) effectors is fundamental for uncovering the mechanisms of pathogenicity and variability, thereby paving the way for developing durable and effective control strategies for stripe rust. However, due to the lack of an efficient genetic transformation system in *Pst*, progress in effector function studies has been slow. Here, we modeled the structures of 15,201 effectors from twelve *Pst* races or isolates, a *Puccinia striiformis* isolate, and one *Puccinia striiformis* f. sp. *hordei* isolate using AlphaFold2. Of these, 8,102 folds were successfully predicted, and we performed sequence- and structure-based annotations of these effectors. These effectors were classified into 410 structure clusters and 1,005 sequence clusters. Sequence lengths varied widely, with a concentration between 101-250 amino acids, and motif analysis revealed the presence of known effector motifs such as [Y/F/W]xC and RxLR. Subcellular localization predictions indicated a predominant cytoplasmic localization, with notable chloroplast and nuclear presence. Clear annotations based on sequence and structure included superoxide dismutase and trehalose-6-phosphate phosphatase. A common feature observed was the formation of similar structures from different sequences. In our study, one of the comparative structural analyses revealed a new structure family with a core structure of four helices, including Pst27791, PstGSRE4, and PstSIE1, which target key wheat immune pathway proteins, impacting the host immune function. Further comparative structural analysis showed similarities between *Pst* effectors and effectors from other pathogens such as AvrSr35, AvrSr50, Zt-KP4-1, and MoHrip2, highlighting convergent evolutionary strategies. This comprehensive analysis provides novel insights into *Pst* effectors’ structural and functional characterization, advancing our understanding of *Pst* pathogenicity and evolution.

**Author Summary:** Stripe rust, caused by the fungus *Puccinia striiformis* f. sp. *tritici* (*Pst*), is a major threat to wheat crops worldwide. The fungus uses special proteins, called effectors, to bypass the plant’s immune defenses and establish infection. To better understand how these effectors work, we used a computational tool, AlphaFold2, to predict the structures of over 15,000 *Pst* effector proteins. Interestingly, some of the effectors resemble proteins found in other plant pathogens, suggesting that different fungi may evolve similar ways to infect plants. Our research offers new insights into the infection strategies of *Pst* and could lead to new methods for protecting wheat from stripe rust.

## Introduction

Pathogens have evolved in different ways to evade multiple host defense mechanisms and subvert cellular signaling pathways to facilitate infestation, expansion, and colonization. One tactic is to secrete effector proteins into the host in a spatio-temporally controlled manner. These effectors perform various roles in the apoplast or within the plant cell, such as enhancing pathogen entry, undermining the plant immune system, and altering metabolism [1]. Understanding how effector proteins function in their host is crucial for comprehending the interactions between plants and pathogens [2]. However, fungal effector proteins often lack known functional domains, making them hard to identify. It is challenging to predict their roles based on sequence alone since these proteins have very diverse sequences; rapidly evolving or recently emerged, exhibiting a wide range of variations. This diversity and lack of similarity to known proteins make it difficult to pinpoint potential effector candidates and understand their biological functions.

Although the sequence similarity of effectors is low, it is found that effectors have relatively conservative structures and form structural families [3–5]. The WY domain, a common structural motif in oomycete RXLR effectors, features a conserved α-helical fold stabilized by a hydrophobic core, typically containing Trp (W) and Tyr (Y), as demonstrated by the structural elucidation of two sequence-unrelated *Phytophthora* effectors, *Phytophthora capsici* AVR3a11 and *Phytophthora infestans* PexRD2 [6]. Recently, variants known as LWY domains were noticed, e.g. the *Phytophthora* effector PSR2 has a WY domain and other six variants of WY domains [7]. The ToxA-like structural family encompasses effector proteins such as AvrL567-A and AvrL567-D from *Melampsora lini* [8,9] and Avr2 (SIX3) and SIX8 from *Fusarium oxysporum* [10,11], characterized by their structural similarity to ToxA from *Pyrenophora tritici-repentis* [12]. MAX (*Magnaporthe* AVRs and ToxB-like) effectors, featuring a typical six-stranded β-sandwich fold, are crucial for the virulence of *Magnaporthe oryzae* (e.g. AvrPiz-t, AVR1-CO39, AVR-Pia) [13–15] and *Pyrenophora tritici-repentis* (e.g. ToxB) [16], despite their sequence divergence. RALPH (RNase-Like Proteins associated with Haustoria) effectors (e.g. BEC1054) [17], characterized by their RNase-like structure and found predominantly in *Blumeria* fungal species, constitute a notable portion of predicted effectors, showcasing a distinct evolutionary expansion within powdery mildews despite highly divergent sequences [18]. AvrLm4-7 [19] and AvrLm5-9 from *Leptosphaeria maculans* and Ecp11-1 from *Cladosporium fulvum* (now *Fulvia fulva*), despite showing low sequence identity, share the *Leptosphaeria* Avirulence and Suppressing (LARS) fold with candidate effectors predicted in at least 13 different fungi [20]. The *Fusarium oxysporum* f. sp. *lycopersici* (*Fol*) dual-domain (FOLD) effectors, exemplified by Avr1 (SIX4) and Avr3 (SIX1), represent a newly identified structural class of effectors with two distinct domains [11]. However, due to the few effectors uncovered by structural biology, the counts are approximately 70 for bacteria, 20 for oomycetes, and 80 for fungi (S1 Table), the structural family of effector proteins found is limited. In recent years, based on TrRosetta, AlphaFold2 and other AI tools to predict protein structure and carry out structural classification, it has been found that there are effectors with low sequence similarity among pathogens but whose structural similarity can be classified into the above said known and novel effector structure family [21–27].

Stripe (yellow) rust, caused by *Puccinia striiformis* f. sp. *tritici* (*Pst*), is a serious fungal disease affecting wheat production areas worldwide, posing a significant threat to global food security [28]. Introgressive hybridization breeding for yellow rust resistance (*Yr*) genes is the most effective, environmentally sustainable, and cost-effective strategy to control stripe rust disease [29]. However, with the rapid evolution of new races overcoming specific resistance genes and emerging *Pst* virulence, wheat varieties often lose their resistance in a short period [28]. The rapid variation of *Pst* virulence may be related to its rich effectors and the variability of their subcellular locations within the wheat cell. Genome sequences of multiple *Pst* races or isolates have been analyzed, predicting about 1,000 to 2,000 secretome or effectors for each race [30]. Despite this, since 2011, only about 50 *Pst* effectors (S2 Table) have been identified experimentally [31,32]. Few effectors have been analyzed for their function because of the obligate biotrophic nature of the pathogen requiring a plant host and a lack of an efficient, reliable, and stable transformation system of the urediniospores, making it challenging to study the mechanism of each effector through genetic methods. Nevertheless, the structural analysis assists in determining the function, i.e. Pst_13661 is the only effector protein with a determined structure enabling a combatting strategy for *Pst* [33,34]. The effector that can be specifically recognized by the host nucleotide-binding leucine-rich repeat receptor (NLR) and cause an immune response is the protein encoded by the avirulence gene (Avr). The failure of wheat varieties carrying NLR-type Yr genes to resist *Pst* may be related to Avr mutations, which render the Yr protein unrecognizable and unable to trigger immunity. Currently, no Avr of *Pst* has been identified. However, five Avr genes have been identified in *Puccinia graminis* f. sp. *tritici* (*Pgt*), a close relative of *Pst*, the causal agent of stem rust [35]. These Avr genes, AvrSr50 [36,37], AvrSr35 [38–40], AvrSr22 [41], AvrSr13 [41], and AvrSr27 [42,43], have been confirmed to be recognized by their corresponding wheat NLRs: Sr50, Sr35, Sr22, Sr13, and Sr27. Recently, it has been reported that the structure of pathogen-secreted proteins can be predicted using AlphaFold2 or other tools and annotated by protein structure databases such as PDB, CATH, and SCOP [23,44,45]. One reason for the slow progress in identifying *Pst* effectors is that predicted effector sequences often lack functional annotations and domain information on protein annotation websites, making it difficult to understand their molecular mechanisms. So far, no study has been reported focusing on large-scale structure predictions or structural annotations of *Pst* effectors.

In this study, by analyzing twelve *Pst* races and isolates, one *Puccinia striiformis* f. sp. *hordei* isolate, and one *Puccinia striiformis* isolate, which consists of 21 protein sets in total, we collected 357,396 proteins and predicted 15,201 effector proteins based on their sequences. Of these, 8,102 had high-confidence predicted structural folds, resulting in the identification of 410 structure clusters and 1,005 sequence clusters. Among the 8,102 effectors, 20.9% have sequence annotations, 75.4% have structure annotations, 6.7% are sequence-related, and 44.2% are structurally related to identified *Pst* effectors. In addition to conventional methods of effector characterization, this structural annotation approach can significantly enhance the efficiency and comprehensiveness of effector analysis. Remarkably, we discovered AvrSr35-like, AvrSr50-like, Zt-KP4-1-like, and MoHrip2-like *Pst* effector candidates with little or no sequence similarity, yet they exhibited conserved structural features. Understanding the structure and relationships of *Pst* effector proteins enhances our insight into their biological functions. This knowledge will be crucial in unraveling the pathogenic mechanisms of wheat stripe rust and in developing new control strategies.

## Results

### Structure prediction of *Pst* effector candidates with AF2

357,396 proteins were collected from 21 proteomes within 14 *Puccinia striiformis* races or isolates: 12 from *Puccinia striiformis* f. sp. *tritici* (*Pst*), 1 from *Puccinia striiformis* isolate 11-281, and 1 from *Puccinia striiformis* f. sp. *hordei* 93TX-2 (S3 Table). We predicted the proteins with signal peptide as secretomes using SignalP 6.0 and excluded proteins annotated with transmembrane domains in InterPro or predicted to have a glycosyl-phosphatidyl-inositol (GPI)-anchor by NetGPI, resulting in the prediction of 27,444 secreted proteins. Further prediction with EffectorP 3.0, followed by redundancy removal across different races or isolates, identified 15,201 effector candidates. Structural predictions for the mature sequence (excluding the signal peptide) of these effector candidates were performed using AlphaFold2 (AF2) (Fig 1A). Following AF2 modeling for 15,201 predicted effectors, nearly half could not be reliably modeled across the 14 races or isolates (Fig 1B and 1C). This likely reflects the high specificity of these effectors, making it difficult to match them to sufficient template structures during AF2 modeling. We filtered out structures with pTM (predicted Templated Modeling) scores < 0.5 and pLDDT (predicted Local Distance Difference Test) scores < 70, retaining high-confidence models. This process yielded 8,102 effectors with reliable structural predictions across 14 races or isolates (S3 Table). We then performed clustering, annotation, and comparative analysis of these 8,102 effectors from both sequence and structural perspectives for further investigation (Fig 1A). Among the 8,102 effector candidates, sequence lengths ranged from 43 to 939 amino acids, with a concentration of 2,221 effectors within the length 101-250 amino acids, consistent with the typical feature of effector lengths (Fig 1D).

**Fig 1.**
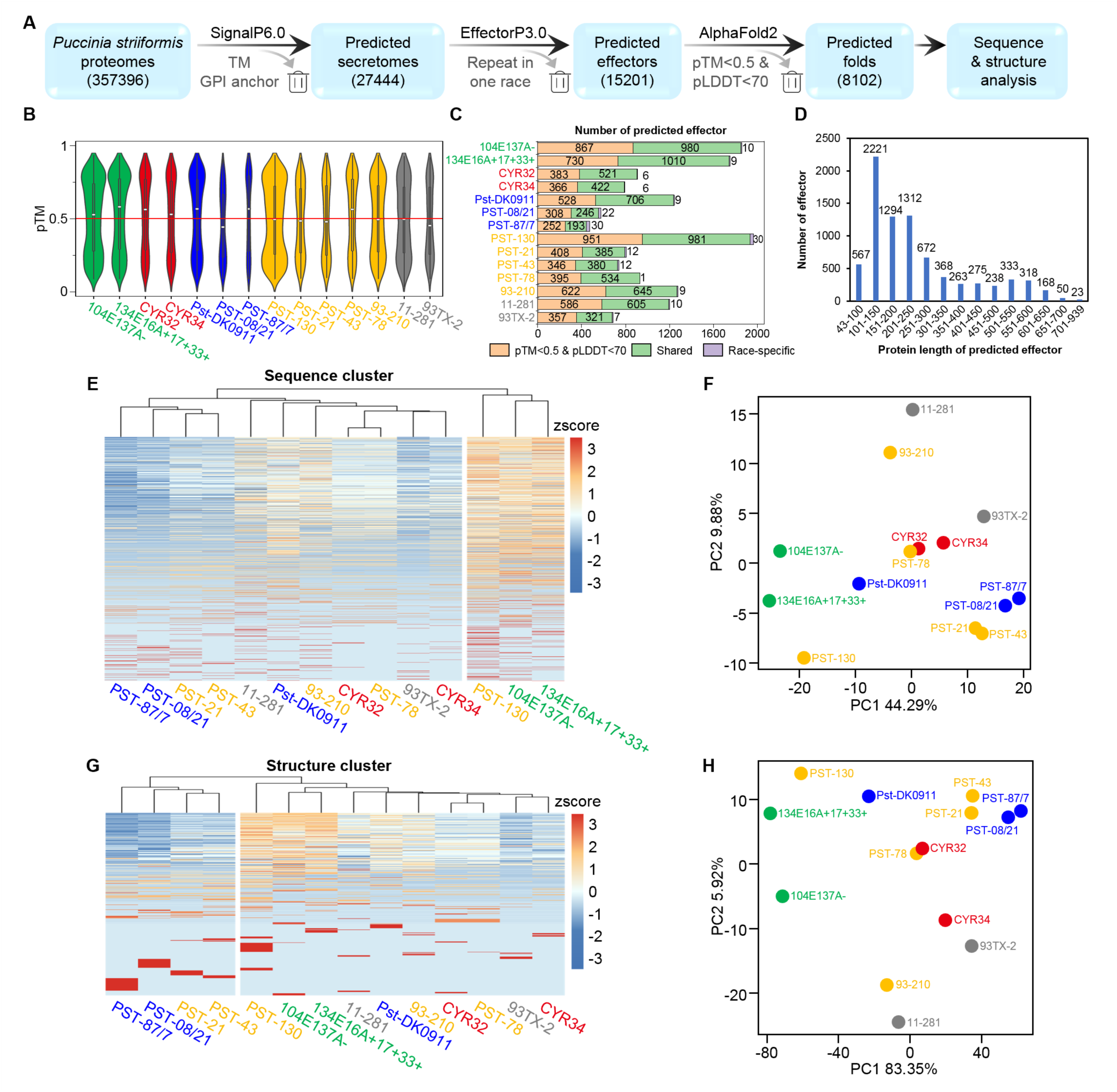
Effector prediction pipeline and statistical analysis of prediction and clustering. (**A**) Bioinformatics workflow for predicting and analyzing effectors in *Puccinia striiformis* (*Ps*) proteomes. Blue boxes denote prediction steps with the number of retained proteins in brackets. Analytical tools are above the arrow; filtered steps are below. Out of 15,201 predicted effectors, 8,102 with reliable structures predicted for further investigation. (**B**) Distribution of pTM scores for evaluating the quality of structure predictions among the 15,201 predicted effectors. The box plot within the violin plot shows the interquartile range (25th to 75th percentiles), with the median represented by a white dot. The whiskers extend to the minimum and maximum values, capped at 1.5 times the interquartile range. The red line represents a pTM value of 0.5; structures with pTM scores below 0.5 were removed. (**C**) Number of 15,201 predicted effectors categorized in 14 *Ps* races or isolates. Orange indicates proteins that did not fold well, green denotes ‘shared’ effectors, and purple denotes ‘race-specific’ effectors. ‘Shared’ effectors have structure cluster members from multiple races or isolates, while ‘race-specific’ effectors do not. (**D**) Sequence length distribution of the 8,102 effectors with reliable structure predictions. (**E, F**) Cluster heatmap and principal component analysis (PCA) showing the number of effectors from each cluster of 1,005 sequence clusters assigned within 14 *Ps* races or isolates. (**G, H**) Cluster heatmap and PCA showing the number of effectors from each cluster of 410 structure clusters assigned within 14 *Ps* races or isolates. *Pst* races or isolates are colored based on their region of first discovery: green for Australia, red for China, blue for Denmark and the UK, yellow for the US, and grey for *Puccinia striiformis* isolate 11-281 and *Puccinia striiformis* f. sp. *hordei* isolate 93TX-2. The same color scheme is applied in Figs. 4-6.

### Clustering of *Pst* effector candidates with sequence and structural comparison

We performed sequence clustering on the mature sequences of 8,102 effector candidates with high-confidence structural predictions using CD-HIT with a threshold of 0.5, resulting in 1,005 sequence clusters (S4 Table). Additionally, we used Foldseek clustering for predicted structures of 8,102 effector candidates with a threshold of 0.5, resulting in 410 structure clusters (S4 Table). Clusters are ordered according to the number of effectors they contain, with Structure Cluster No. 1 (Struc.C_1) having the most effector candidates. Of these 410 structure clusters, 165 structure clusters were single-protein clusters. 7,929 effectors were classified as shared, and 173 effectors were classified as race-specific (Fig 1C). We analyzed the distribution of 8,102 effectors within sequence and structure clusters across 14 races or isolates using cluster heatmaps and principal component analysis (PCA). In general, the quantitative characteristics and classification relationships of effectors in 14 races or isolates exhibit consistency in both sequence and structure clusters, suggesting that effectors with similar sequences tend to form similar structures. Notably, there is a close correspondence between the sequence clusters and structure clusters of effectors from CYR32 and PST-78, indicating similarity in effector components between these two *Pst* races. Furthermore, the relationships between sequence clusters and structure clusters of effectors from PST-87/7, PST-08/21, PST-21, and PST-43 show similarities. Interestingly, despite having different hosts, CYR34 and 93TX-2 share similarities in terms of both sequence and structure for their effectors (Fig 1E-1H).

### A comprehensive understanding of *Pst* effector candidates’ characterization with sequence-based and structural annotation

Cysteine richness is a characteristic feature of effectors. We analyzed the cysteine content in the mature sequences of the 8,102 effectors. Among these, 1,970 effectors contain 6 cysteine residues. Remarkably, 4 effectors have 30 cysteine residues, 13 of which have 31 cysteine residues (Fig 2A) with an average length of 458 amino acids. These 17 effectors belong to Struc.C_27 and Sequence Cluster No. 62 (Seq.C_62). They are predicted to be apoplastic effectors but also contain nuclear localization signals (S4 Table). Additionally, many short effectors (<100 amino acids) have a higher cysteine content, such as effectors belonging to Struc.C_27 and Seq.C_62, the average length of mature sequences is 55 amino acids and contain 8 cysteines, with a cysteine content of 14.5% (S4 Table).

**Fig 2.**
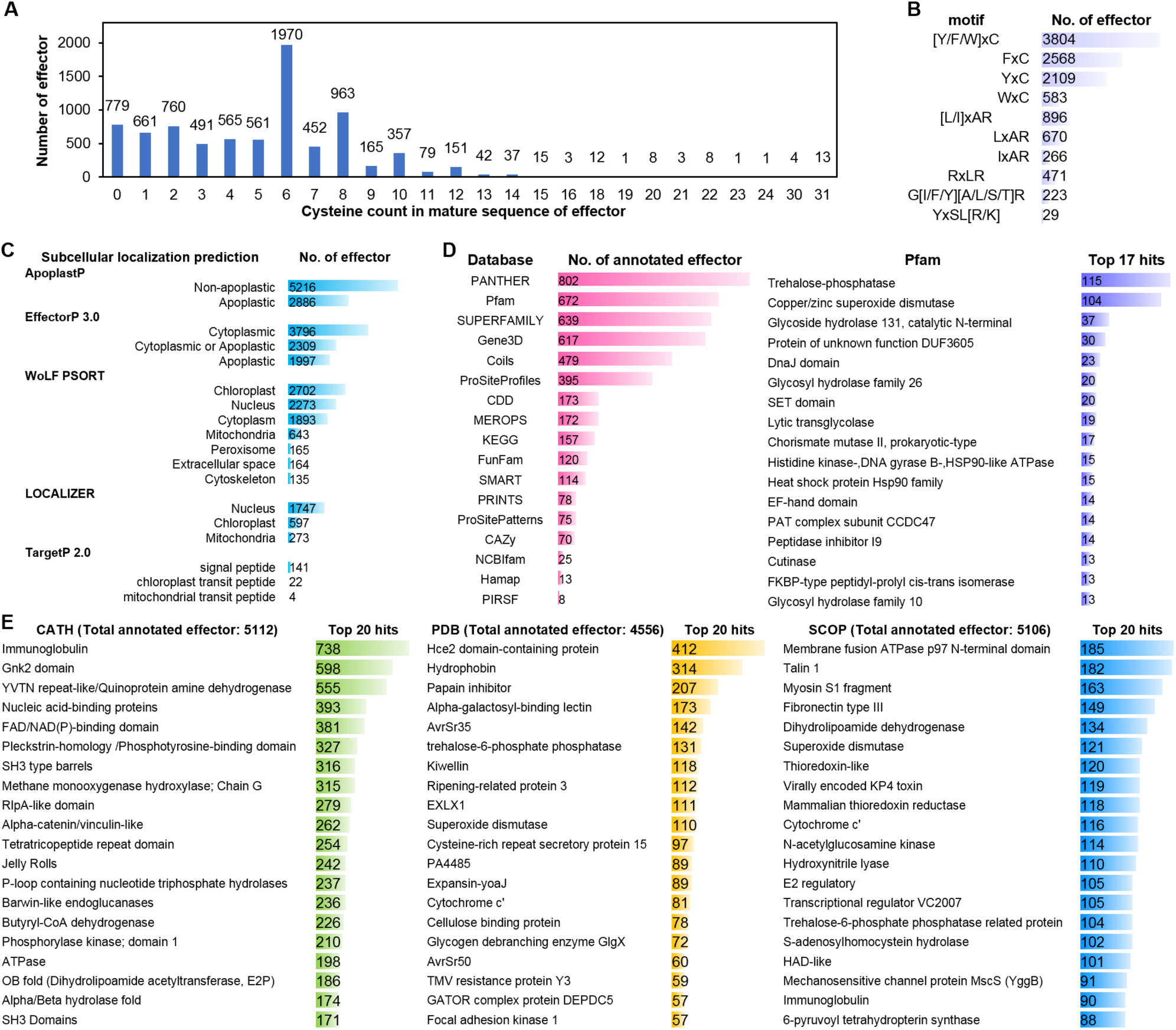
Statistical analysis of effector characteristics. (**A**) Statistics of cysteine count in the mature sequences of effectors. (**B**) Number of motif-containing effectors. (**C**) Number of effectors in different subcellular localization predictions. (**D**) Number of effectors in various protein sequence annotation databases, and statistics of the top 17 hits of effectors annotated within Pfam. (**E**) Statistics of the top 17 hits of effectors structurally annotated within CATH, PDB, and SCOP.

Classical effector motif analysis (Fig 2B and S4 Table) revealed that 3,804 effectors (approximately 47% among 8,102 effector candidates) contain the [Y/F/W]xC motif, which is found in the wheat powdery mildew and rust effector candidates, predominantly in the forms of FxC and WxC. Additionally, 896 effectors possess the [L/I]xAR motif, which is found in some effectors of *Magnaporthe oryzae*, with LxAR being the most common. The RxLR motif, a common effector motif from oomycetes and fungi, is present in 471 effectors. Another 223 effectors contain the G[I/F/Y][A/L/S/T]R motif, which is found in some effectors of *Melampsora lini*. A total of 29 effectors have the YxSL[R/K] motif, which is found in oomycetes. We checked motif [R/K]CxxCx12H which is found in *Magnaporthe oryzae* and [R/K]VY[L/I]R, which is found in *Blumeria graminis* f. sp. *hordei*, but not detected. This indicates that different pathogens have their own specific motif characteristics.

We used ApoplastP, EffectorP 3.0, LOCALIZER, WoLF PSORT, and TargetP 2.0 for subcellular localization prediction of the 8,102 effectors (Fig 2C and S4 Table). Although the predictions varied across different programs, the overall trend indicated that more effectors were predicted to be cytoplasmic rather than apoplastic, with primary localizations in the chloroplast and nucleus. Notably, 141 effectors’ mature sequences were predicted to have signal peptides by TargetP 2.0, but SignalP 6.0 did not detect any signal peptides in these mature effector sequences. Therefore, when predicting subcellular localization using different programs, it is essential to conduct a comprehensive and integrated analysis to ensure the accuracy of the predictions.

We annotated the sequences of effector candidates using various protein databases, including PANTHER, Pfam, SUPERFAMILY, Gene3D, Coils, ProSiteProfiles, CDD, MEROPS, KEGG, FunFam, SMART, PRINTS, ProSitePatterns, CAZy, NCBIfam, Hamap, and PIRSF (Fig 2D and S4 Table). Overall, 1,695 effectors (approximately 21% among 8,102 effector candidates) had sequence annotation information. Specifically, 802 effectors were annotated by PANTHER, 672 by Pfam, 639 by SUPERFAMILY, and 617 by Gene3D. Among all protein databases, Pfam provided the most annotation entries (19k entries, accessed on July 18, 2023). Since Pfam includes the most comprehensive and numerous annotations, we examined the top statistics from Pfam to know sequence annotation of effector candidates in general. The top three Pfam annotations were trehalose-phosphatase, copper/zinc superoxide dismutase, and glycoside hydrolase 131 catalytic N-terminal.

In addition to sequence-based annotation, we utilized the predicted structures of effector candidates using AF2 and compared them with protein structure annotation information from CATH (Protein Structure Classification Database; Class, Architecture, Topology, Homologous superfamily), PDB (Protein Data Bank), and SCOP (Structural Classification of Proteins) by Foldseek. This approach provided structural annotation information for 6,110 effectors (approximately 75% among 8,102 effector candidates), significantly more than sequence-based annotations (S4 Table). Specifically, 5,112 effectors were annotated by CATH, 4,556 by PDB, and 5,106 by SCOP. We analyzed the top 20 hits for each database. The most frequently occurring annotations were immunoglobulin in CATH, Hce2 domain-containing protein (or named as *Zymoseptoria tritici* effector Zt-KP4-1) in PDB, and membrane fusion ATPase p97 N-terminal domain in SCOP (Fig 2E). Although the annotation methods of CATH, PDB, and SCOP differ and no identical annotations appear in the top 20 hits of all three databases, certain annotations were common between PDB and SCOP, such as superoxide dismutase, cytochrome c’, and trehalose-6-phosphate phosphatase. Superoxide dismutase and trehalose-phosphatase are also the major annotations in sequence annotation. These annotations may represent prominent features of *Pst* effectors.

### *Pst* effector candidates reflect progressively differentiating structure

From the 8,102 effectors, we selected 1,178 representative effectors (marked in grey in S4 Table) from distinct structure and sequence clusters to perform a pair-wise TM-align analysis using Foldseek by filtering out pairs with TM scores < 0.5. This resulted in a structure cluster network graph with 1,015 nodes and 32,119 edges. We present the representative structures of the top ten largest structure clusters (Fig 3A). The largest structure cluster is Struc.C_1 containing 683 effectors. Struc.C_2 to Struc.C_5 contains 438, 326, 323, and 254 effectors, respectively. The top ten structure clusters together account for 36.6% of the effectors, representing the overall characteristics of *Pst* effector candidates. By observing the structure cluster network, we can identify the expansion and variation patterns of effector structures. For example, the structures of Struc.C_10 and Struc.C_1 are quite similar, but Struc.C_10 forms multiple tandem repeats of the Struc.C_1 structure. Similarly, Struc.C_3 appears as a double tandem structure of Struc.C_9. Struc.C_2 shows a dispersed expansion trend, potentially indicating faster structural variation and the gradual formation of new effector clusters like Struc.C_7. The effector structures of Struc.C_4, Struc.C_6, and Struc.C_8 are similar but have gradually diverged structurally. In contrast, Struc.C_5 has a relatively simple structure with fewer associations with other major structure clusters.

**Fig 3.**
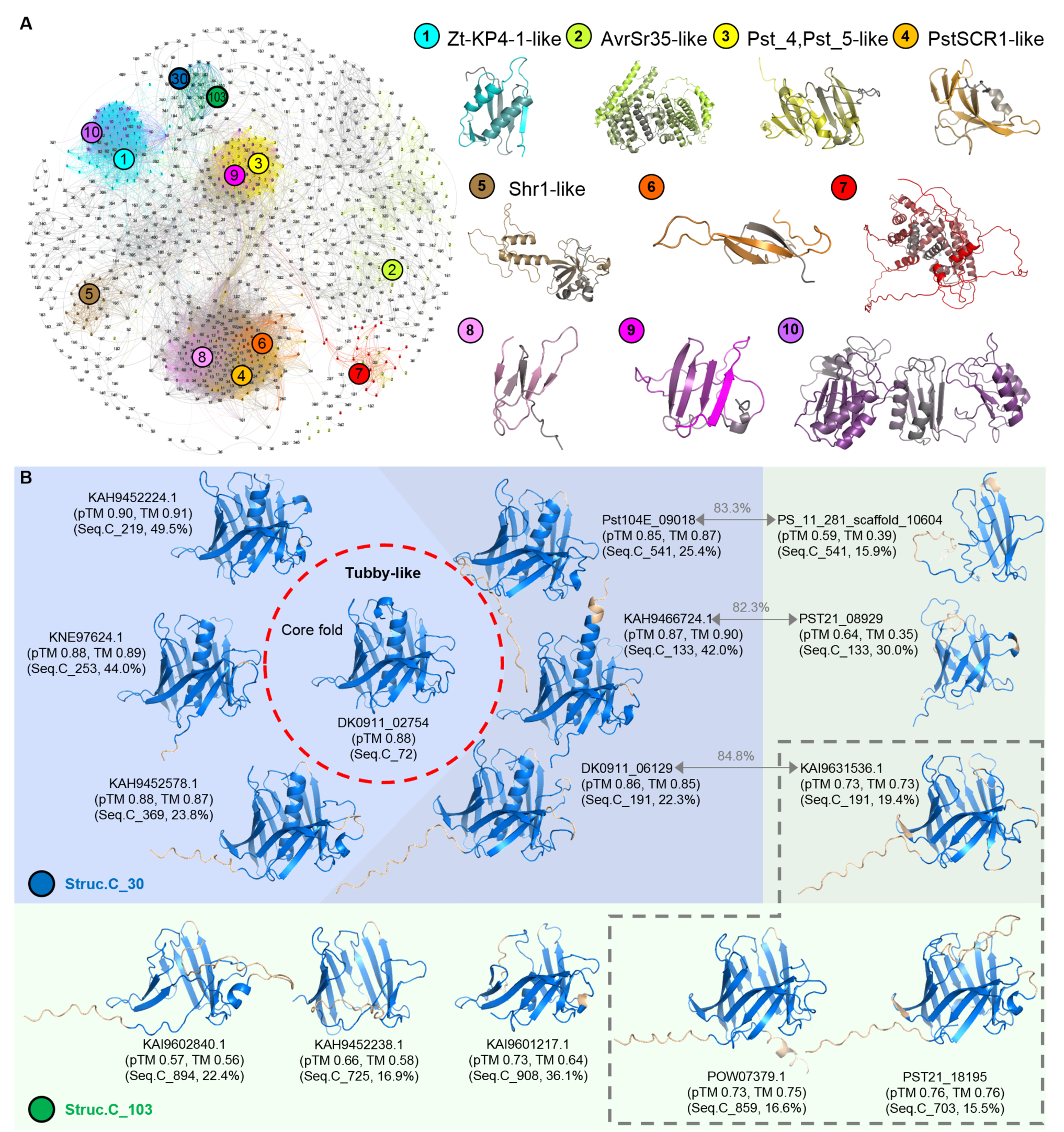
Relationship of *Pst* effector structure clusters. (**A**) Structural similarity network of representative effectors from each structure cluster’s sequence cluster (1,178 proteins marked in grey in S4 Table). The 10 major structure clusters, along with structure clusters No. 30 (Struc.C_30) and No. 103 (Struc.C_103), are highlighted. Representative structures from the 10 major structure clusters are shown. (**B**) Structures from Struc.C_30 are shown with a blue background, and structures from Struc.C_103 are shown with a green background. The effector DK0911_02754, highlighted within the red dashed circle, is annotated as Tubby-like based on its sequence, with its predicted structure used as the core fold. Parts of other displayed structures that do not superimpose with the core fold are marked in light orange. The transparent grey background indicates three pairs of effectors from the same sequence cluster but belonging to Struc.C_30 and Struc.C_103, respectively. The sequence similarity between these pairs is indicated above the connecting arrows. Effectors within the grey dashed box are from Struc.C_103 and lack the central α-helix found in Struc.C_30 members. The pTM values from AF2 modeling, TM-scores (TM) compared with the core fold, sequence cluster numbers (Seq.C), and sequence identity with the core fold sequence are shown below the protein IDs.

### The structure of *Pst* effector candidates is more conserved than the sequence

Our analysis of the correspondence between structure and sequence clusters (S5 Table) reveals that effectors within the same sequence cluster are predominantly distributed into a single structure cluster. This indicates that similar sequences tend to form similar structures as observed in the cluster heatmap analysis (Fig 1E and 1G). However, a single structure cluster can encompass multiple sequence clusters. For instance, Struc.C_2 includes 61 sequence clusters, Struc.C_1 contains 52 sequence clusters, and Struc.C_3 comprises 41 sequence clusters (S5 Table). This suggests that different effector sequences can also form similar structural conformations.

Taking Struc.C_30 as an example, it includes 7 sequence clusters, with protein sequence similarities between clusters being less than 50%, some even less than 25% (Fig 3B). Notably, only some effectors from Seq.C_72 have sequence annotation from SUPERFAMILY, identifying them as a Tubby C-terminal domain-like. The Tubby-like domain is characterized by a β-barrel structure enclosing an internalized α-helix in the center of the barrel. The predicted structure of Seq.C_72’s effector in Struc.C_30, such as DK0911_02754, matches this structural feature. The remaining 6 sequence clusters within Struc.C_30 also exhibit high pTM scores. Furthermore, the representative proteins of these 6 sequence clusters show high structural similarity to the representative protein of Seq.C_72 (DK0911_02754) based on US-align analysis, indicating that they are all Tubby-like domain proteins despite some having low sequence similarity (22%-25%) with Seq.C_72 (Fig 3B) illustrating different effector sequences can form similar structures.

Further investigation revealed that Seq.C_133, Seq.C_191, and Seq.C_541, which are part of Struc.C_30, are also included in Struc.C_103. Proteins in Struc.C_103 are notably missing the complete central α-helix found in Struc.C_30, and some also lack parts of β-sheet in the β-barrel (Fig 3B). Despite including proteins from different sequence clusters, Struc.C_103 exhibits similar structural characteristics. This observation highlights that different sequences can form similar structural conformations. However, even though proteins from Seq.C_133, Seq.C_191, and Seq.C_541 share over 80% sequence similarity, their structural TM scores are relatively low and they are distributed into different structure clusters. This suggests that these proteins have undergone different evolutionary paths, potentially leading to divergent functions or loss of functional structural features.

### Identified *Pst* effectors represent a new *Pst* effector structural family

To date, over 50 *Pst* effectors have been identified. We performed a BLASTP alignment of these identified effectors against the mature sequences of 8,102 effector candidates, using a threshold of query coverage and percent identity ≥ 50%. This sequence alignment identified 547 effector candidates as sequence homologous hits (S2 Table and S4 Table). We compared the predicted structures of 8,102 effector candidates with the identified *Pst* effectors and *Pgt*-Avr effectors (S2 Table), as well as with PDB chains including Pst_13661 (PDB chain: 8hf9_A), AvrSr27 (PDB chain: 8v1j_A), AvrSr35(PDB chain: 7xx2_B, 7xc2_D, 7xds_A, 7xds_B, 7xe0_B, 7xvg_B) and AvrSr50(QCMJC) (PDB chain: 7mqq_A), using Foldseek and US-align for pair-wise comparison. Structures with TM-scores greater than 0.5 by both Foldseek and US-align were structure homologous hits of the identified *Pst* effectors and *Pgt*-Avr, resulting in 3,707 effectors (S2 Table and S4 Table). The sequence and structural comparisons are summarized in bubble plots (Fig 4A and 4B). Overall, structural comparisons enabled the identification of more effectors than sequence-based comparisons.

**Fig 4.**
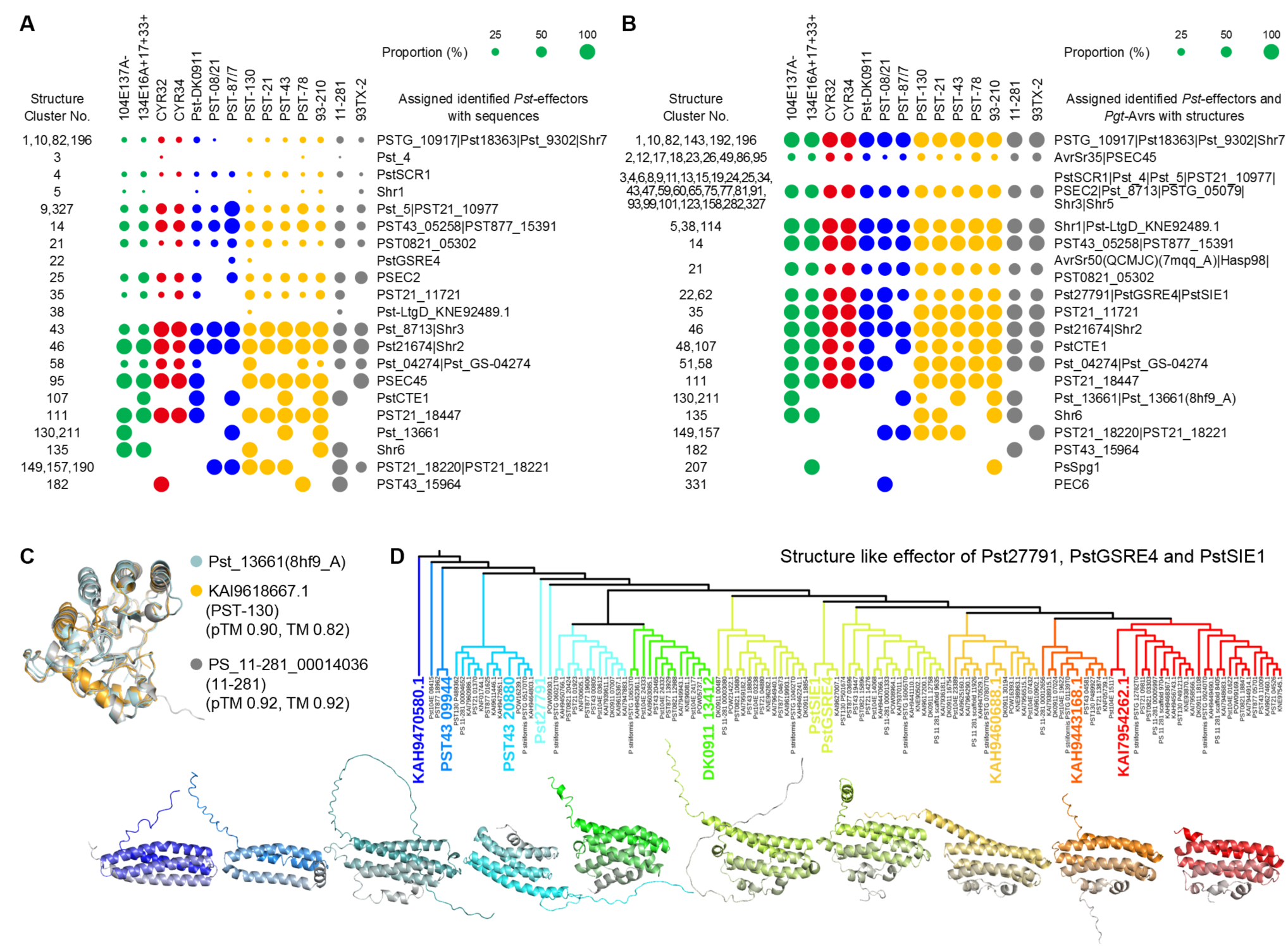
Sequence-based, structural and phylogenetic analysis between identified *Pst* effectors and *Pst* effector candidates. (**A**) Proportion of identified *Pst* effectors assigned to effector candidates from 14 *Ps* races or isolates within structure clusters, based on sequence comparison, shown with circles of varying sizes. (**B**) Proportion of identified *Pst* effectors with predicted structures and PDB structures of Pst_13661 (8HF9_A), AvrSr35 (7XX2_B, 7XC2_D, 7XDS_A, 7XDS_B, 7XE0_B, 7XVG_B), and AvrSr50(QCMJC) (7MQQ_A) assigned to predicted structure of effector candidates from 14 *Ps* races or isolates within structure clusters, shown with circles of varying sizes. (**C**) Pst_13661 (8HF9_A) structure and Pst_13661-like effector predicted structures. The pTM values from AF2 modeling and TM-scores (TM) compared with the Pst_13661 (8HF9_A) structure are indicated below the protein IDs. (**D**) Structural phylogenetic tree analysis of predicted structures of identified *Pst* effectors Pst27791, PstGSRE4, PstSIE1 and effector candidates from Struc.C_22 and Struc.C_62. Representative structures of groups are shown in the same color as their groups in the phylogenetic tree, with structures displayed from N-terminus (colored) to C-terminus (grey).

Pst_13661 is currently the only *Pst* effector with a resolved structure (Liu *et al*., 2023). Based on the sequence of Pst_13661, homologous sequences were found in the races 104E137A-, PST-87/7, PST-78, and 93-210 (Fig 4A). In addition to the four aforementioned races, structurally similar homologs were also identified in *Pst* race PST-130 and *Ps* isolate 11-281 (Fig 4B). The predicted structures of the homologs in *Pst* race PST-130 and *Ps* isolate 11-281 are highly accurate and show significant similarity to the structure of Pst_13661 (PDB chain: 8hf9_A) (Fig 4B and 4C).

Interestingly, based on sequence similarity, PstGSRE4 homologs were found only in the effectors of *Pst* races PST-87/7 and PST-130, which belong to Struc.C_22 (Fig 4A). However, based on structural similarity, PstGSRE4, along with two other effectors, Pst27791 and PstSIE1, which do not share sequence similarity, showed predicted structural similarity to approximately 71% of the effectors in races or isolates belonging to Struc.C_22 and Struc.C_62 (Fig 4B). We then conducted a structural phylogenetic analysis of 125 effectors from 12 sequence clusters within Struc.C_22 and Struc.C_62, as well as the three identified effectors PstGSRE4, Pst27791, and PstSIE1 using DALI (Fig 4D). This analysis assembled them into nine groups. Among these, the core structure of the second group, consisting of Pst104E_08415, PST877_18962, and PST43_09944, was formed by three helices, while the other groups had a core structure of four helices. Previous studies have reported that the host wheat interactors for Pst27791, PstGSRE4, and PstSIE1 are TaRaf46 [46], TaCZSOD2 [47], TaGAPDH2 [48], and TaSGT1 [49], respectively, which are key proteins in wheat immune pathways. This led us to identify a class of widely present effectors in *Pst* with a core structure of four helices that commonly interact with key immune pathway proteins during the infection of wheat.

### The structure of *Pst* effector candidates from multiple different sequence clusters is similar to AvrSr35 and AvrSr50

Previous research predicted several *Pst* Avr candidates, including 48 secreted proteins and 14 non-secreted proteins [50]. We compared the sequences and the predicted structures of these 62 *Pst* Avr candidates with the sequences and predicted structures of 8,102 effector candidates (Fig 5A, S4 Table and S6 Table). None of the non-secreted *Pst* Avr candidates showed sequence or predicted structural similarity to any of the effector candidates supporting our prediction that the effectors originate from the secretome. To date, no *Pst* Avr has been cloned and identified. However, in the closely related wheat rust pathogen *Puccinia graminis* f. sp. *tritici* (*Pgt*), five Avrs have been cloned and identified, they are AvrSr50, AvrSr35, AvrSr22, AvrSr13, AvrSr27. None of these *Pgt* Avrs showed sequence similarity to the 8,102 effectors candidates, AvrSr22, AvrSr13, and AvrSr27 nor did they show structural similarity. Notably, AvrSr35 and AvrSr50 exhibited structural similarity to several effectors with different sequences (Fig 5B and 5C, S2 Table and S4 Table). Among them, effector candidates distributed in Struc.C_2, Struc.C_12, Struc.C_17, Struc.C_18, and Struc.C_26 from 26 sequence clusters are structurally AvrSr35-like, with a concentration in Struc.C_2. Effector candidates distributed in Struc.C_21 from 6 sequence clusters are structurally AvrSr50-like.

**Fig 5.**
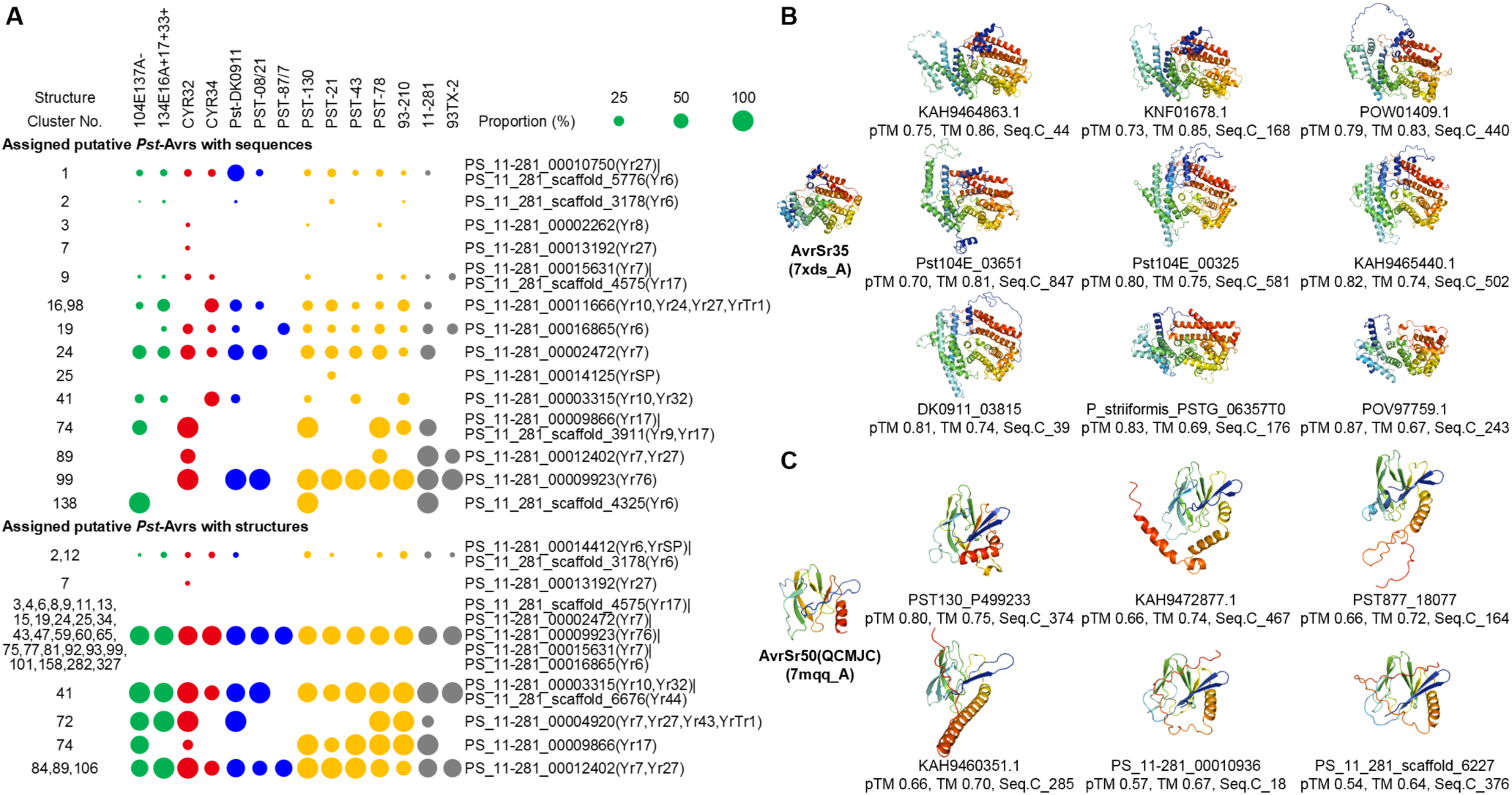
Sequence-based and structural analysis of putative *Pst* Avr candidates. (**A**) Proportion of putative *Pst* Avr candidates assigned to effector candidates from 14 *Ps* races or isolates within structure clusters, based on sequence and predicted structure comparison, shown with circles of varying sizes. (**B, C**) Structural analysis of AvrSr35 (7XDS_A) and AvrSr50(QCMJC) (7MQQ_A), along with predicted structures of AvrSr35-like and AvrSr50-like *Pst* effectors from different sequence clusters. The pTM values from AF2 modeling, TM-scores (TM) compared with AvrSr35 (7XDS_A) or AvrSr50(QCMJC) (7MQQ_A) structures, and sequence cluster numbers (Seq.C) are indicated below the protein IDs. Structures are shown from N-terminus (blue) to C-terminus (red).

### *Pst* effector candidates are structurally similar to some effectors from bacteria, oomycetes, and other fungi

In addition to *Pst* and *Pgt*, the structures of approximately 180 effectors from various bacteria, oomycetes, and fungi have been determined (S1 Table). None of these effectors showed sequence similarity to the 8,102 effector candidates using BlastP, indicating the distant phylogenetic relationships between these pathogens and *Pst*. However, some of these effectors exhibited structural similarities to the *Pst* effector candidates (Fig 6A, S1 Table and S4 Table).

**Fig 6.**
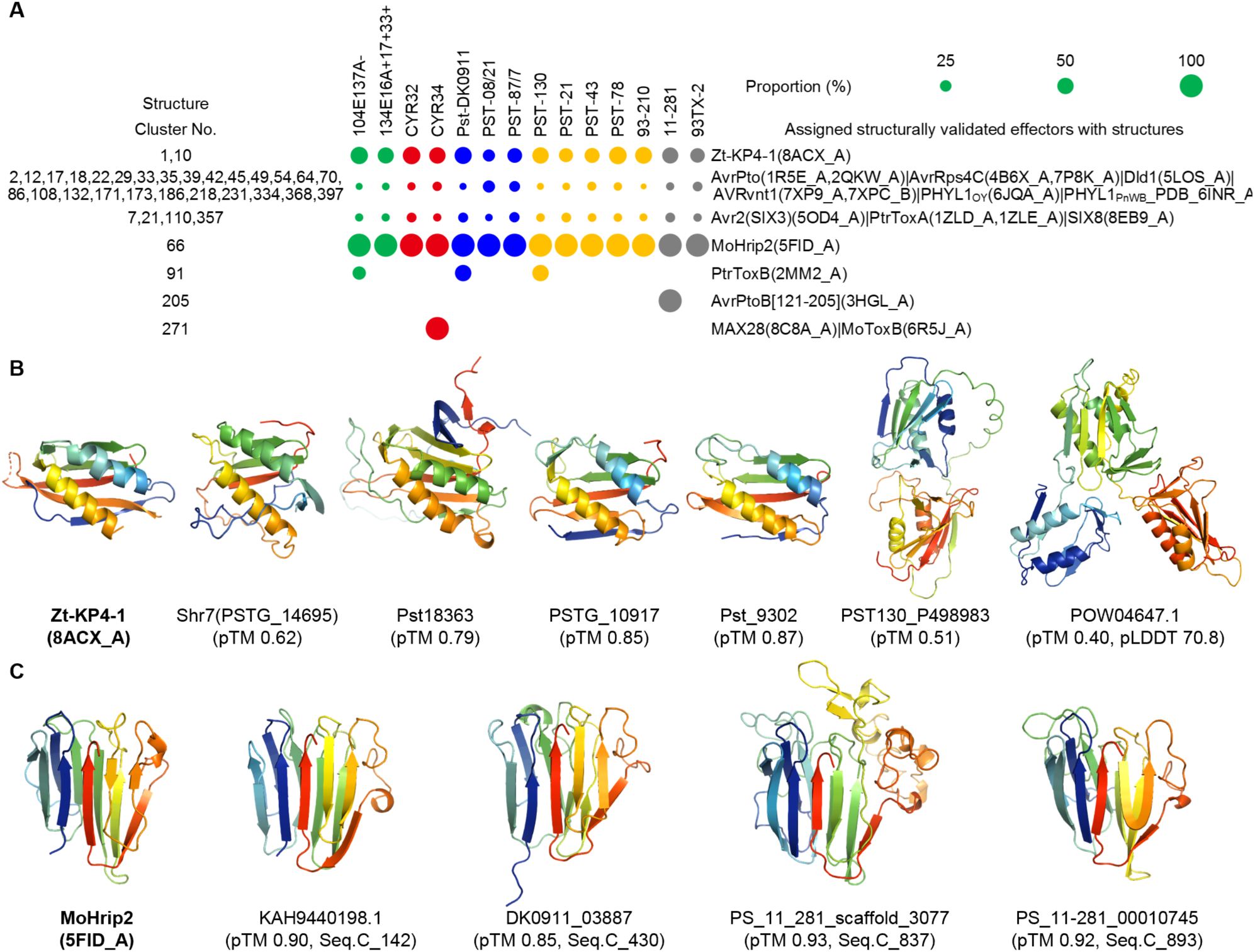
Structural analysis between structurally validated effectors and *Pst* effector candidates. (**A**) Proportion of structurally validated effectors assigned to the predicted structure of effector candidates from 14 *Ps* races or isolates within structure clusters, shown with circles of varying sizes. (**B, C**) Structural analysis of Zt-KP4-1 (8ACX_A) and MoHrip2 (5FID_A), along with representative predicted structures of Zt-KP4-1-like and MoHrip2-like *Pst* effectors. The pTM values from AF2 modeling are indicated below the protein IDs. For MoHrip2-like effectors, the sequence cluster number (Seq.C) is also indicated. Structures are shown from N-terminus (blue) to C-terminus (red).

Several effectors characterized by two helices, such as AvrPto, AvrRps4C, Dld1, AVRvnt1, PHYL1_OY_, and PHYL1_PnWB_, showed structural similarity to 14% of the effectors distributed across various structure clusters from all races or isolates. Additionally, 15% of the effectors from all races or isolates distributed in Struc.C_7, Struc.C_21, Struc.C_110, and Struc.C_357 exhibited structural similarities to effectors from the Tox-like structure effector family, such as Avr2 (SIX3), PtrToxA, and SIX8. Some effectors from Struc.C_91 and Struc.C_271 also showed structural similarity to effectors from the MAX structure effector family, including PtrToxB, MAX28, and MoToxB.

Interestingly, Zt-KP4-1 showed structural similarity to 48% of the effectors distributed in Struc.C_1 and Struc.C_10 from all races or isolates. Among the effectors in Struc.C_1 and Struc.C_10 with structural similarity to Zt-KP4-1, some also exhibited sequence or structural similarity to identified *Pst* effectors Shr7 (PSTG_146695), Pst18363, PSTG_10917, and Pst_9302. Zt-KP4-1 indeed shared structural similarity with these identified *Pst* effectors (Fig 6B). Notably, Struc.C_1 contains the highest number of effectors in this study, suggesting that Zt-KP4-1-like structures are widespread in *Pst*. Additionally, we observed effectors such as PST130_P498983 with a tandem Zt-KP4-1-like structure and POW04647.1 with a triplet Zt-KP4-1-like structure (Fig 6B), indicating an expansion in the evolution of *Pst* effectors. Most significantly, MoHrip2 showed high structural similarity to all effectors in Struc.C_66 from all races or isolates and was exclusively similar to effectors in Struc.C_66 (Fig 6A and 6C).

## Discussion

Our comprehensive and detailed analysis of *Pst* effector candidates sheds light on the landscape of characterizations and the evolutionary dynamics of *Pst* effector repertoire, offering new insights into the molecular basis of stripe rust pathogenesis and informing future strategies for disease management. We have collected a large number of *Pst* effector candidates for sequence and structural annotation, providing an essential resource for subsequent *Pst* effector research. This study is the first to perform structural clustering of *Pst* effector candidates examining their relationships from a structural perspective. We have also explored and discovered for the first time, from a structural viewpoint, the existence of effector structural families among *Pst* effector candidates and their structural similarities with known effectors from other pathogens. This research offers valuable insights for the study of effectors of different pathogens, demonstrating how large-scale sequence and structural analyses can elucidate effector characteristics and advance effector research. Previous reviews or articles have summarized effectors with known structures [2,23]. Building on this foundation, we have further enhanced and supplemented the data, ultimately identifying over 180 structures (S1 Table). These findings will serve as a crucial training dataset for future research on effector structure, structural prediction, and structural family classification.

### AlphaFold2 facilitates effector research

Although this study focused on the sequence and structural analysis of 8,102 *Pst* effector candidates with well-predicted structures, it is important to note that nearly half of the predicted effectors were not included in this research due to poor structural predictions (pTM scores < 0.5 and pLDDT < 70). One possible reason for this is the strong sequence specificity of these *Pst* effectors, resulting in fewer reference templates during AF2 model construction, which hampers accurate modeling. However, this suggests that these effectors may have a higher specificity to *Pst*, potentially making them key effectors in the successful infection of wheat. For effectors with reference models, such as Pst_13661-like effectors, reliable models with pTM scores above 0.79 were obtained, demonstrating the potential for high-confidence predictions. Despite the capabilities of AF2 as a protein prediction tool, structural biology experiments remain essential for analyzing highly specific effectors, expanding the number of template models, and increasing prediction accuracy. AF2, while being a leading AI protein structure prediction tool, also has limitations. It cannot account for the cellular content where protein functions, such as pH, salt concentration, ions, and post-translational modifications, which are critical for protein conformation [51]. The newer AlphaFold3 has addressed some of these issues, such as introducing ions and ligands in structural predictions. Although there is substantial room for improvement in AF2, it has significantly advanced large-scale research on effector proteins based on their structures. For example, understanding the structure of effectors can aid in the design of compounds as effector inhibitors [34]. AF2 can also facilitate protein interaction studies [52,53], greatly aiding in the exploration of pathogen-host interactions through the study of effector-interactor mechanisms [54].

### Structural annotations assist effector characterizations

Before conducting effector cloning and functional identification, researchers often use protein sequence functional annotation databases to preliminarily predict the biological functions of effectors and obtain corresponding research direction ideas. These predictions are often unsatisfactory. In this study of 8,102 *Pst* effector candidates, even after searching through 17 protein sequence functional or domain annotation databases, only about 21% of the effectors had functional or domain annotations based on their protein sequences. However, when we compared the predicted structures of *Pst* effector candidates with the PDB, CATH, and SCOP databases, and filtered out structural annotations with TM-scores < 0.5, approximately 75% of the effector candidates still had structural annotation information. In this way, structure-guided similarity searches have made it possible to better annotate effector repertoires. Since structure often determines function, structural information can further predict or infer interactors within the host. The databases comparable in Foldseek are constantly being updated allowing comparisons not only with PDB, CATH, and SCOP but also with other databases to obtain more annotation information. Therefore, predicting the structures of *Pst* effectors and subsequently comparing them with protein structure annotation databases to obtain annotation information will provide preliminary ideas for the early stages of effector identification research. Our findings provide a detailed understanding of the structural and functional complexity of *Pst* effectors, offering a foundation for future studies on effector biology and host-pathogen interactions. Experimental validation and functional assays of these predicted effectors will be essential to fully understand their roles in *Pst* pathogenicity and host resistance.

### Diverse sequences of effectors share a structural commonality

From the analysis of the relationship between the sequence and structure of effectors (Fig 1E and 1G, S5 Table), generally, effectors with similar sequences can form similar structures, aligning with our expectations. However, there are still instances where effectors with similar sequences form structures with low similarity. For example, in (Fig 3B), effectors belonging to the same sequence cluster with at least 82% sequence similarity form different structures and thus belong to different structure clusters. Therefore, in addition to using sequences to identify homologs, structural predictions can also aid in determining homologs. This study also found that effectors with different sequences can form similar structures. There are 129 structure clusters, which are ‘sequence-unrelated structurally similar’ (SUSS) clusters among 410 structure clusters, found in this study. This may arise from the pathogen using amino acid resources optimally during the infection process to form functionally similar effectors with similar structures.

### New *Pst* effector families found

Although *Pst* cannot be cultured, and there is a lack of an efficient and reliable system for stable transformation, making it challenging to study its pathogenicity mechanisms through genetic methods, over 50 *Pst* effectors have already been identified (S2 Table). Previous research often studied effectors as isolated entities with little integration among them. So far, it has only been discovered that the identified *Pst* effectors Pst_12806, Pst_4, and Pst_5 share the same host wheat interactor TaISP [55,56] and that there is an interaction between PstCEP1 and PSTG_11208 [57]. In this study, by comparing the sequences and predicted structures of identified *Pst* effectors with 8,102 *Pst* effector candidates, we found that many identified *Pst* effectors have a widespread sequence or structural homologs in different races or isolates. Furthermore, we discovered a more typical class of *Pst* effectors with a core structure of four helices, represented by the identified *Pst* effectors Pst27791, PstGSRE4, and PstSIE1, forming a unique *Pst* effector structural family. This finding provides a new perspective for future effector research, suggesting that by studying effector structural families comprehensively, we can better understand how these structural families of effectors function in host infection. It may be interesting to test the silencing of the effectors having highly similar structures, in addition to silencing the alleles to determine the function of any given effector. We have also observed that Struc.C_7, which contains 206 effector candidates, lacks annotation information and does not exhibit sequence or structural similarity to any of the identified effectors. However, as a structure cluster of effectors widely present in *Pst*, it remains uncharacterized and requires further study.

### Potential *Pst* Avr candidates and convergent evolutionary strategies

Identifying *Pst* Avr genes is crucial for understanding *Pst* variability. Although there have been reports predicting *Pst* Avr candidates, no *Pst* Avr has been identified to date. In this study, by comparing the predicted structures of *Pst* effector candidates with those of *Pgt* Avr, we identified many effectors that are AvrSr35-like and AvrSr50-like. These AvrSr35-like and AvrSr50-like *Pst* effector candidates could potentially be cognate Avr candidates for particular yellow rust resistance proteins and can be further experimentally validated. We also discovered that pathogens employ convergent evolutionary strategies for their effectors. Specifically, we found that the predicted structures of *Pst* effector candidates show high structural similarity to effectors from bacteria, oomycetes, and other fungi, despite having no sequence similarity in BlastP comparisons. In this study, at least 5.3% of the effector candidates are Zt-KP4-1-like and are primarily distributed in Struc.C_1 and Struc.C_10, indicating a commonality among *Pst* effectors.

The widespread presence of Zt-KP4-1-like structures among *Pst* effectors points to potential conserved mechanisms in effector evolution and function. Furthermore, the identification of structural similarities with known effectors from other pathogens, despite low sequence similarity, highlights the importance of structural analysis in uncovering functional relationships. These similarities suggest that effectors from different pathogens might converge on similar host targets or pathways, offering potential cross-species insights into effector biology.

## Materials and Methods

### Collection of *Puccinia striiformis* proteome and identified effectors and putative *Pst* Avr candidates

Proteomes of 14 races or isolates for *Puccinia striiformis* consisting of 357,396 proteins were collected (S3 Table). 54 identified *Pst* effectors, 5 avirulence factors (Avr) from stem rust and 4 unpublished *Pst* effectors were collected from the literature and our lab, respectively (S2 Table). The 181 known effector structures were downloaded from PDB (S1 Table) (https://www.rcsb.org/, downloaded on 2024-06-13). 62 putative *Pst* Avrs i.e. 48 secreted and 14 non-secreted (S6 Table) were collected from Li et al., 2020 [50].

### Effector prediction

SignalP 6.0 was used to identify the secreted proteins [58], glycosyl-phosphatidyl-inositol (GPI) anchoring containing proteins excluded with the help of NetGPI 1.1 [59]. The remaining candidates excluded with the InterProscan 5.63-95.0 if they have transmembrane in it or if their signal peptides overlapped with PFAM domains over ten or more amino acids [60]. To determine the effectors, EffectorP 3.0 was used for effector prediction, including their cytoplasmic or apoplastic localization in the host [61].

### Motifs analyses and subcellular localization prediction

Common effector motifs from oomycetes and fungi were searched, including RxLR [62], and YxSL[R/K] detected in oomycetes [63], [L/I]xAR and [R/K]CxxCx12H in some effectors of *Magnaporthe oryzae* [64], [R/K]VY[L/I]R identified in *Blumeria graminis* f. sp. *hordei* [65], [Y/F/W]xC found in the wheat powdery mildew [66] and rust effector candidates and G[I/F/Y][A/L/S/T]R in some effectors of *Melampsora lini* [67]. ApoplastP [68], LOCALIZER [69], TargetP 2.0 [70], and WoLF PSORT [71] were used to predict the subcellular localization of effectors.

### Effector characterization

Full-length sequences of effectors were used to search in the InterProscan 5.63–95.0 [60] against all databases available i.e CDD, Coils, FunFam, Gene3D, Hamap, NCBIfam, PANTHER, Pfam, PIRSF, PRINTS, ProSite Patterns, ProSite Profiles, SMART, SUPER FAMILY. To find the cysteine residue count, mature sequences of effectors were used. MEROPS was used to find the peptidases, for this purpose HMMER was employed with the MEROPS database against our sequences (https://www.ebi.ac.uk/Tools/hmmer/search/phmmer, accessed on 2024–4–25). CAZy (Carbohydrate– Active Enzymes Database) term annotation performed with mature effector sequences in eggNOG– mapper 2.1 (accessed on 2024-4-24) [72]. KEGG Orthology Search was conducted on KofamKOALA (accessed on 2024-4-24) [73].

### Structure prediction

The structures of 17,158 putative *Pst* effectors, using their mature sequences, were predicted by AlphaFold2 via the LocalColabFold approach [74,75]. Additionally, the structures of identified effectors, avirulence factors of stem rust, and putative *Pst* Avr candidates (S1 Table, S2 Table, and S6 Table) were also predicted using the same method to efficiently utilize our resources. The computational workload for LocalColabFold was efficiently managed using the Lenovo ThinkBook 16p Gen 4, equipped with an Nvidia RTX4090 graphics card and the 13th Gen Intel™ Core i9 processor. Five models were generated, and the ranked_1 model was chosen as it had the best pLDDT score. All structures were then filtered based on pLDDT and pTM scores, retaining those with a pLDDT of 70 or above and/or a pTM of 0.5 or above for further analysis.

### Clusters analyses

To create sequence clusters, CD-HIT was used with the sequence identity threshold set to 0.5 [76]. For structure clustering, the Foldseek cluster option was used with a 0.5 threshold to group similar structures into the same cluster [77].

### Structural annotation

We used Foldseek [77] for structural similarity search against the PDB chain, SCOPe40 and CATH50 databases [78–80], downloaded from Foldseek on 22 March 2024. Homologues with a TM score greater than 0.5 were retained, with a maximum of ten hits per query.

### Network analysis

We selected 1,170 representatives from our dataset (marked in grey in S4 Table) and computed their pair-wise TM scores using Foldseek. Those edge with TM scores > 0.5 were then imported into Gephi 0.10.1 with layout of Fruchterman-Reingold to construct and visualize our network for further analysis.

### Homologous effectors search

We used Foldseek and US-align [81] tools to compare predicted protein structures from various sources i.e 54 identified *Pst* effectors, 5 Avrs from stem rust, 4 unpublished *Pst* effectors, 62 putative *Pst*–Avrs, and 181 effector structures downloaded from the PDB. We identified matches with a TM score of 0.5 or higher as potentially homologous. We utilized BlastP [82] to identify sequence similarities among the proteins mentioned above, considered matches where the query coverage and percent identity were both 50% or higher.

### Program and software

We used TBtools–II [83] for converting FASTA files to tables and running InterProScan analyses in a loop. Origin 2022 facilitated the creation of violin plots, histograms, and stacked bar charts. DALI was utilized to generate Newick dendrograms for structural comparisons [84]. iTOL (https://itol.embl.de/) was used for visualizing phylogenetic tree analyses. Cluster heatmap and PCA (principal component analysis) were performed using the OECloud tools at https://cloud.oebiotech.com. Protein structures were visualized and edited in the PyMOL [85].

## Acknowledgements

We would like to thank Chaoming Zhang for his assistance in predicting the protein structures for specific races using AlphaFold2. We also extend our gratitude to Diane G. O. Saunders and Cristobal Uauy for providing the proteome data of PST-21, PST-43, PST-87/7, and PST-08/21. Their contributions were invaluable to our research.

## Supporting information

**S1 Table. Identified and structurally resolved effectors or avirulence factors of plant pathogens with their structure homologous analyses.**

**S2 Table. Identified *Puccinia striiformis* f. sp. *tritici* effectors and avirulence factors of *Puccinia graminis* f. sp. *tritici* with their homologous analyses.**

**S3 Table. Protein sets of 14 *Puccinia striiformis* races or isolates used for analyses and statistics from proteome to effectors.**

**S4 Table. The metadata for 8,102 predicted effectors with well-fold from 14 *Puccinia striiformis* races or isolates.**

**S5 Table. Statistics between sequence cluster and structure cluster of *Puccinia striiformis* f. sp. *tritici* effector candidates.**

**S6 Table. Putative avirulence factors of *Puccinia striiformis* f. sp. *tritici* with their homologous analyses.**

## Author Contributions

**Conceptualization:** Mahinur Akkaya

**Data Curation:** Yulei Wang, Nan Wu, Raheel Asghar

**Formal Analysis:** Raheel Asghar, Nan Wu

**Investigation:** Raheel Asghar, Nan Wu

**Methodology:** Raheel Asghar, Nan Wu, Mahinur Akkaya

**Resources:** Mahinur Akkaya

**Software:** Noman Ali, Nan Wu, Raheel Asghar

**Supervision:** Mahinur Akkaya

**Validation:** Nan Wu, Mahinur Akkaya

**Writing – Original Draft Preparation:** Mahinur Akkaya Raheel Asghar, Nan Wu

**Writing – Review & Editing:** Raheel Asghar, Nan Wu, Mahinur Akkaya, Noman Ali, Yulei Wang

## Data Availability

15,201 predicted effectors fasta file, 8,102 AF2 predicted structures of *Pst* effector candidates, and AF2 predicted structures of identified *Pst* effectors, *Pgt* Avrs and *Pst* Avr candidates can be downloaded from Zenodo (https://doi.org/10.5281/zenodo.12946213).

